# An engineered Abcb4 expressing model reveals the central role of NF-κB in the regulation of drug resistance in zebrafish

**DOI:** 10.1101/2020.03.30.016824

**Authors:** Cong-Jie Sun, Rong-Yin Hu, Zhi-Cao Li, Lu Jin, Chuan Ye, He Lu, Yan-Hua Zhou, Ting Zhou, Zhi-Xu He, Li-Ping Shu

## Abstract

Multidrug resistance (MDR) represents the major cause of unsatisfaction in the application of chemotherapy for cancer treatment. So far, an *in vivo* robust high-throughput screening system for anti-tumor drug MDR is still lacking and the molecular mechanisms for MDR still remain elusive. Given a myriad of merits of zebrafish relative to other animal models, we aimed to establish MDR system in zebrafish stably expressing ATP-binding cassette (ATP-cassette) superfamily transporters and study the potential regulatory mechanism. We first constructed a *Tg(abcb4:EGFP)* transgenic zebrafish stably expressing both Abcb4 and EGFP using Tol2-mediated approach. The expression level of Abcb4 and EGFP was significantly induced when *Tg(abcb4:EGFP)* transgenic zebrafish embryos were exposed to doxorubicin (DOX) or vincristine (VCR), accompany with a marked decrease in rhodamine B (RhB) accumulation in embryos, which indicates a remarkable increase in drug efflux upon the exposure to DOX or VCR. Mechanistically, AKT and ERK signaling were activated when treated with DOX or VCR. With the application of AKT and ERK inhibitors, the drug resistance phenomena could be reversed with differential responsive effects. Of note, downstream NF-κB played a central role in the regulation of Abcb4-mediated drug resistance. Taken together, the engineered *Tg(abcb4:EGFP)* transgenic zebrafish model provides a new platform for drug resistance screening *in vivo*, which could facilitate and accelerate the process of drug development.

## Introduction

To date, great efforts have been made to advance cancer treatments and clinically significant progress has been made to improve therapeutic outcomes for patients. However, drug resistance is still a major problem in cancer therapy, leading to ineffective anticancer treatments via intrinsic or acquired mechanism [1, 2]. As cancer is the second leading cause of death after cardiovascular disease, drug resistance is responsible for 90% of all cancer related deaths [3-5]. Thus, it stresses the need not only to better understand the underlying mechanism of drug resistance, but also to establish robust and practical experimental model systems for detecting and screening drug resistance.

Despite of numerous of studies detailing the characteristics of drug resistance, it still remain unsolved which justifies the complicated mechanisms. Among these mechanisms, multi-drug resistance (MDR) was drawn particular attention, which appears to be the major impediment to the success in cancer chemotherapy, due to the overexpression of ATP-binding cassette (ABC) family of drug efflux transporters, especially P-glycoprotein (P-gp; ABCB1) [6]; and drug efflux transporters was shown to be expressed in many clinical cancers, effectively excluding many clinically cancer chemotherapy drugs, including doxorubicin, vincristine, paclitaxel and etoposide, contributing to drug resistance and therapeutic failure [7-9]. In particular, compelling evidence has shown the correlation between the expression of ABCB1 and the chemotherapy sensitivity and survival rate of cancer patients [10-13]. These studies highlight the key role of ABCB1 in MDR, rendering ABCB1 as an important indicator for monitoring MDR.

Given the notorious role of drug resistance in cancer therapy, it absolutely requires a robust and practical model to determine drug resistance. As such, zebrafish has come under the spotlight, given a myriad of advantages relative to other animal models, in particular the gene manipulation. Meanwhile, the application of zebrafish model in the high-throughput screening of small molecules has also become a rapid and effective approach for the study of specific biological processes, especially in the research on highly conserved developmental regulation, disease genes and signal transduction pathways [14-16]. As such, we aimed to engineer a transgenic zebrafish model expressing drug efflux transporter P-gp promoter driven fluorescent reporter gene by using Tol2-meidated approach and investigate the potential regulatory mechanism. We believe that this model will provide a robust platform for drug resistance screening *in vivo*.

## Materials and Methods

### Chemicals

Doxorubicin hydrochloride (CAS#: 25316-40-9), gefitinib (CAS#: 184475-35-2), rhodamine B (CAS#: 81-88-9), and vincristine sulfate (CAS#: 57-22-7) were purchased from Sigma-Aldrich (Shanghai, China). U0126 and LY294002 were bought from Cell Signaling Technology (Danvers, MA, USA). BAY 11-7082 was obtained from Abcam (Cambridge, MA, USA). RPMI-1640 medium was sourced from Gibco (Gibco, Thermo Scientific, IL, USA). T4 DNA Ligase and JetFlex™ Genomic DNA Purification Kit was obtained from Invitrogen (Thermo Scientific, IL, America). KOD DNA polymerase was purchased from Toyobo. (Osaka, JPN) RhB was dissolved in MilliQ water and the rest of chemicals were prepared in dimethyl sulfoxide (DMSO, Sigma-Aldrich). Final concentration of DMSO in exposure medium is at 0.1%.

### Zebrafish strain and fish husbandry

The 3-month-old adult zebrafish was housed and maintained in our laboratory as previously described [17]. Embryos were obtained from natural spawning, and then collected and incubated in medium containing 19.3 mM NaCl, 0.23 mM KCl, 0.13 mM MgSO_4_ 7H_2_O, 0.2 mM Ca (NO_3_)_2_, and 1.67 mM Hepes (pH 7.2). The protocol was approved by the Animal Experimentation Committee of Guiyang Medical University, Guizhou, People’s republic of China.

### Cell line and cell culture

Human myelogenous leukemia line K562 was obtained from the American Type Culture Collection (ATCC). The cells were maintained in suspension in RPMI-1640 medium supplemented with 10% FBS at 37°C in a humidified atmosphere of 5% CO2.

### Genomic DNA purification

Genomic DNA was purified from 100 embryos from 24, 48, 72, 96, and 120 hpf zebrafish embryos using the JetFlex™ Genomic DNA Purification Kit according to the manufacturer’s instructions. The quality of extracted genomic DNA was assessed using gel electrophoreses and the concentration was measured by Nanodrop (ND-1000; Thermo Scientific). The resultant genomic DNA was stored at −20□°C until further analysis.

### Cloning of the zebrafish *abcb4* promotor

The promoter region −838/+694 of *abcb4* was amplified by genomic PCR with KOD DNA polymerase (the primers listed in Table 1). The promoter fragments were then inserted into the pGL3-Enhancer vector between the *Nhe*I and *Xho*I sites with T4 DNA Ligase. The resultant recombinants were transformed into Escherichia coli DH5α and the construct was confirmed by DNA sequencing.

**Table 1.**
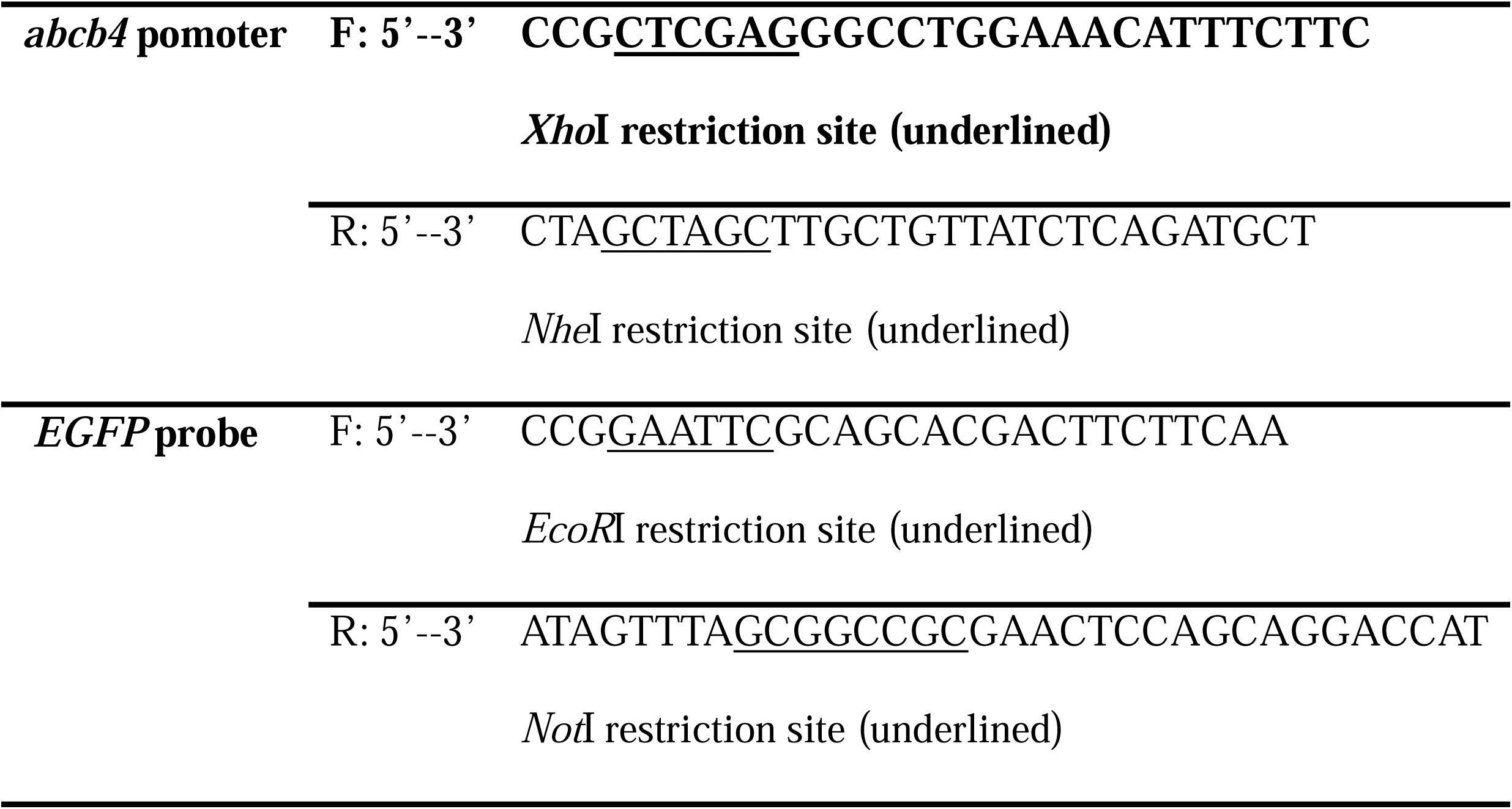
The sequences for the primers.

### Dual-luciferase reporter assay

Promoter activity was detected using the dual-luciferase reporter assay system (Promega, Madison, WI, USA) according to the manufacturer’s instructions. Briefly, K562 cells were transfected with 0.1 μg of Renilla luciferase expression plasmid pRL-TK (internal control for normalizing transfection efficiency; Promega) and 0.4 μg of *abcb4* promoter construct, pGL3-control plasmid (positive control; Promega), or pGL3-enhancer plasmid (negative control; Promega). The firefly luciferase readings were normalized by the Renilla luciferase reading to calculate the relative fold-change.

### Micro-injection and generation of stable transgenic zebrafish lines

To generate the *Tg(abcb4:EGFP)* zebrafish line, the *abcb4* promoter and EGFP were inserted into the T2AL200R150G vector between the *Nhe*I and *Xho*I sites with T4 DNA Ligase [18]. Tol2 transposase mRNA was synthesized with SP6 RNA polymerase from a pCS-TP plasmid as described previously [19]. Zebrafish fertilized eggs at the one-cell stage were injected with 1 nl of a mixture containing 250 ng/μl circular DNA of *abcb4*:EGFP plasmid and 250 ng/μl transposase mRNA. The injected embryos were raised and crossed with wild-type counterpart. The integrated Tol2 construct was transmitted to the F_1_ generation and confirmed by genotyping.

### Whole-mount mRNA *in situ* hybridization

Whole-mount in situ hybridization (WISH) assay was performed as described previously [17], and WISH staining was analyzed with the stereomicroscope and imaging software (Nikon, JPN).

### Treatment in zebrafish embryos

Embryos were transferred to individual wells of a 12-well plate (50 embryos/well) and incubated in 2 mL of embryo medium with doxorubicin (15 μM, DOX), vinblastine (25 μM, VCR), and gefitinib (15 μM, GEF) respectively. The concentration was determined by LC_50_ value (48 hours). Exposures were started with 72 hpf embryos and terminated after 48 hours. Besides, we employed three inhibitors including BAY 11-7082 (0.5 μM), LY294002 (6.5 μM), and U0126 (2.5 μM) which targets on NF-κB, AKT, and ERK pathways, respectively. The concentration of inhibitors were determined by LC50 values [20]. The 48 hpf embryos were pre-treated with inhibitors for 2 hr, followed by treatment with DOX, VLB, GEF, GEF+DOX, and GEF+VLB until 120 hpf, respectively. The fluorescence of EGFP was detected by fluorescence microscope (Olympus, JPN).

### Quantification of EGFP fluorescence

For quantification of EGFP fluorescence, 50 embryos per treatment were sonicated in 250 μL of a hypotonic lysis buffer (10 mM KCl, 1.5 mM MgCl2, 10 mM Tris HCl, pH 7.4), the sonicates were briefly centrifuged, 200 μL of the supernatant were transferred to a black 96-well microplate (Corning, Sigma-Aldrich, Madison, WI, USA) and the EGFP fluorescence was measured in a fluorescence plate reader (Synergy H1, BioTek, USA). Triplicates of five treatments along with a solvent control were run per experiment.

### Western blot analysis

Western blotting assay was performed to test the expression level of protein of interest in embryos after treatment. Zebrafish embryos at 120 hpf were homogenized in 500 μL of RIPA buffer containing 50 mM Tris, pH 7.5, 150 mM NaCl, 10 mM EDTA, 1% NP-40, 0.1% SDS and 1 mM PMSF with the addition of phosphatase inhibitors. The lysates were collected and centrifuged at 12,000 ×g for 10 min at 4 °C. Protein concentration was determined using DCTM kit (Bio-Rad, Hercules, CA, USA) and denatured for 10 min with SDS-PAGE sample loading buffer. The samples were subjected to Western blotting assay to test protein expression using primary antibodies of AKT (1:1000), p-AKT (1:1000), ERK (1:2000), p-ERK (1:2000), ABCB4 (1:2000), EGFP (1:2000) and P65 (1:2000), p-P65 (1:2000), and β-actin (1:4000) (Cell Signaling Technology, Danvers, MA, USA).

### Rhodamine B accumulation assay

Up to 10 embryos per mL were incubated in the test solutions with 0.5 μM rhodamine B for one hour at 26°C in the dark, rinsed three times with clean culture water to remove dye from the chorion. For quantification of RhB dye uptake, 50 embryos per treatment were sonicated in 250 μL of a hypotonic lysis buffer (10 mM KCl, 1.5 mM MgCl2, 10 mM Tris HCl, pH 7.4), the sonicates were briefly centrifuged, 200 μL of the supernatant were transferred to a black 96-well microplate (Corning) and the rhodamine B fluorescence was measured at 595 nm (emission)/530 nm (excitation) in a fluorescence plate reader (BioTek). Triplicates of five treatments along with a solvent control were run per experiment. The amount of rhodamine B accumulated in zebrafish embryos was quantified with a rhodamine B standard curve.

### Statistical analysis

All experimental results are expressed as means ± standard deviation (SD) and the data were retrieved from at least three independent assays. Statistical analysis was performed using GraphPad Prism 5 (GraphPad, San Diego, CA, USA). The data were evaluated by one-way ANOVA, and the differences between the means were assessed by using Duncan’s test. The P<0.05 was considered as statistically significant.

## Results

### Cloning of *abcb4* gene promoter of zebrafish

Due to the lack of proper animal model for testing MDR, we first applied zebrafish to establish a MDR system *in vivo* expressing *abcb4* gene and validated the activity. The construct was designed to express the promoter of *abcb4* gene which was located between *adam15* gene and the *abcb4* gene within a 7, 492 bp non-coding region (Figure. 1A). Specifically, the promoter was inserted at upstream of the starting codon of the *abcb4* gene within a −838/+694 region, with a size of 1, 532 bp (Figure 1B). Subsequently, the transcription was validated via Dual-luciferase reporter assay. The inserted promoter was abundantly expressed relative to the negative control, evident from the luciferase activity (Figure 1C). Taken together, the results show that the promoter of *abcb4* gene has been correctly and functionally constructed, which paves the way for the subsequent construction of zebrafish model system.

**Figure 1.**
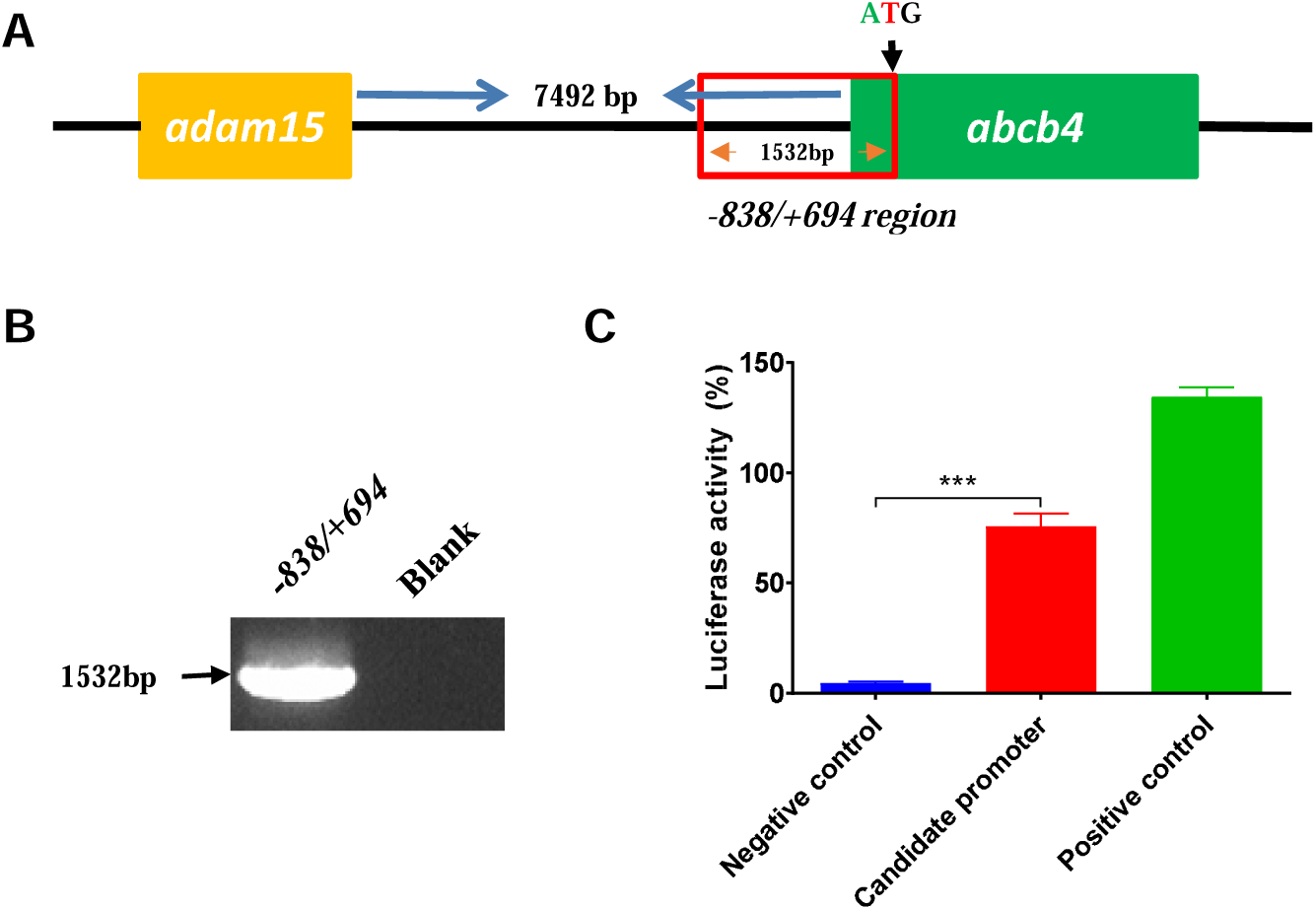
Construction of the reporter vector for the −838/+694 region of the *abcb4* promoter. (A) The diagram of promoter region −838/+694. (B) Gel analysis of the amplification of the −838/+694 region. Lane 1: the amplification of the −838/+694 region detected by 1% agarose GEF; lane 2: blank control. (C) The pGL3 −838/+694 reporter plasmid and pRL-TK were transiently co-transfected into K562 cells. Luciferase activities were measured 48 h after transfection. The results are presented as the relative luciferase activity. Positive control: pGL3-control vector; negative control: pGL3-Enhancer vector. The candidate promoter was compared to the negative control. Data are shown as the means ± SD of at least three independent experiments, \*\*\**P*< 0.001 vs. control group.

### Construction of *Tg (abcb4: EGFP)* zebrafish

Next, the establishment of *Tg (abcb4: EGFP)* zebrafish was performed and verified. The active *abcb4* promoter and *EGFP* were constructed into Tol2 carrier (Figure 2A). The expression was examined via screening 3 generations of transgenic zebrafish, showing a correct transcription *in vivo* (Figure 2B). In subsequence, the genome of F_3_ transgenic zebrafish was sequenced, showing a correct integration of *abcb4* and *EGFP* (Figure 2C). Furthermore, the translation was evaluated via testing the expression of EGFP. The fluorescent signal appeared in intestinal tissue of the 4dpf embryo, to a lesser extent, the cerebral hemisphere. Of note, the fluorescence signal enhanced as time increases (Figure 2D). Additionally, the WISH assay also showed *EGFP* gene expression in the intestinal tissue of the transgenic zebrafish at 4dpf stage (Figure 2E). Collectively, these results show that *Tg (abcb4: EGFP)* transgenic zebrafish model has been established.

**Figure 2.**
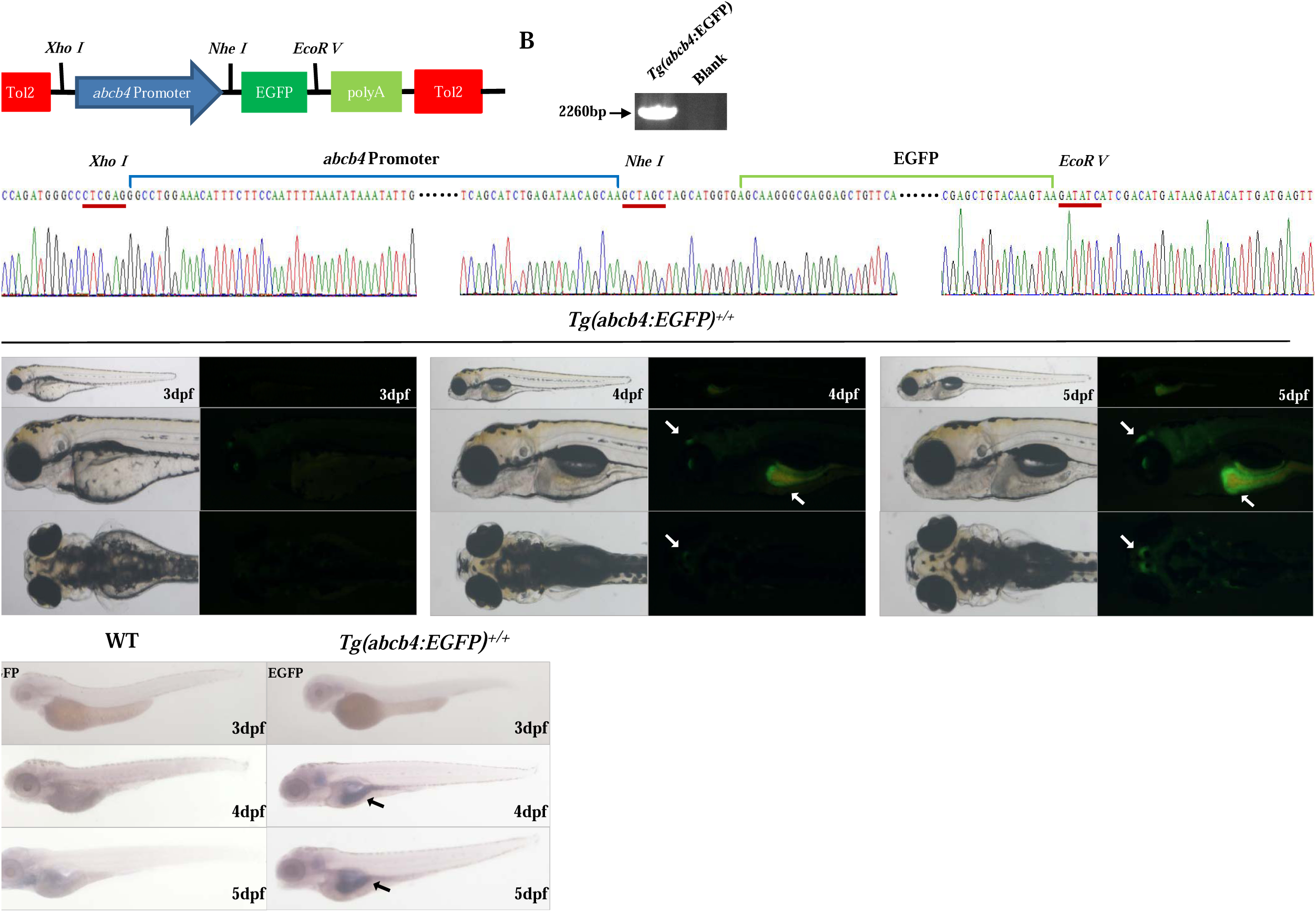
Molecular evidence and expression pattern of EGFP in *Tg(abcb4:EGFP)* zebrafish embryos. (A) Schematic representation of construction in T2AL200R150G vector. (B) Gel analysis of the genomic amplification of the *abcb4:EGFP* fragment for *Tg(abcb4:EGFP)* zebrafish. Lane 1: the genomic amplification of the *abcb4:EGFP* fragment detected by 1% agarose GEF; lane 2: blank control. (C) Sequencing results of the genomic PCR products. (D) Fluorescence micrographs of 3, 4, and 5 dpf *Tg(abcb4:EGFP)* zebrafish embryos. The 4 and 5 dpf embryos express EGFP specifically in the brain and intestine (white arrow). (E) Lateral view of representative 3, 4, and 5 dpf images of zebrafish embryos in which *EGFP* mRNA transcripts where visualized with WISH. In the wild-type line compared with that of the transgenic line, the intestinal bulb and intestine of transgenic zebrafish embryos were strongly stained at 4 and 5 dpf *Tg(abcb4:EGFP)* zebrafish embryos, showing high expression of *EGFP* (black arrow).

### Drug efflux is increased in *Tg(abcb4: EGFP)* transgenic zebrafish

The major challenge of chemotherapy is drug resistance induced therapeutic failure in the clinic, in which patients cannot have long-lasting benefit after the initial responsive effect. Thusly, it requires for better understanding of drug resistance with respect to molecular mechanism and potential targets. Since we have established the Abcb4 expressing transgenic zebrafish model, we first employed this system to test the effect of two conventionally used chemo-cytotoxic drugs (doxorubicin and vincristine) on the expression of Abcb4. It has been reported that administration of doxorubicin and vincristine induces acquired drug resistance without clear mechanism. We exposed the 3dpf *Tg(abcb4: EGFP)* transgenic zebrafish embryos to doxorubicin and vincristine for 48 hours at sub-LC_50_ concentrations (Figure S1), in which the EGFP was significantly increased compared to control zebrafish (Figure 3A and B), which indicates that the expression of Abcb4 was significantly increased after doxorubicin and vincristine exposure. Along with the test of expression level of Abcb4, the function and Abcb4 was also examined. The accumulation of RhB was markedly less than that in control zebrafish embryos, suggesting that the efflux machinery was enhanced in *Tg(abcb4: EGFP)* transgenic zebrafish embryos upon the exposure of doxorubicin and vincristine (Figure 3C). On the other hand, the 3dpf *Tg(abcb4: EGFP)* transgenic zebrafish embryos were treated with gefitinib alone or in combination with doxorubicin and vincristine. In comparison to control embryos, gefitinib did not alter the expression and activity of Abcb4, evident from the EGFP signal and RhB accumulation. Of note, gefitinib normalized the effect of doxorubicin and vincristine on the expression and activity of Abcb4 in *Tg(abcb4: EGFP)* transgenic zebrafish (Figure 3A, B and C).

**Figure 3.**
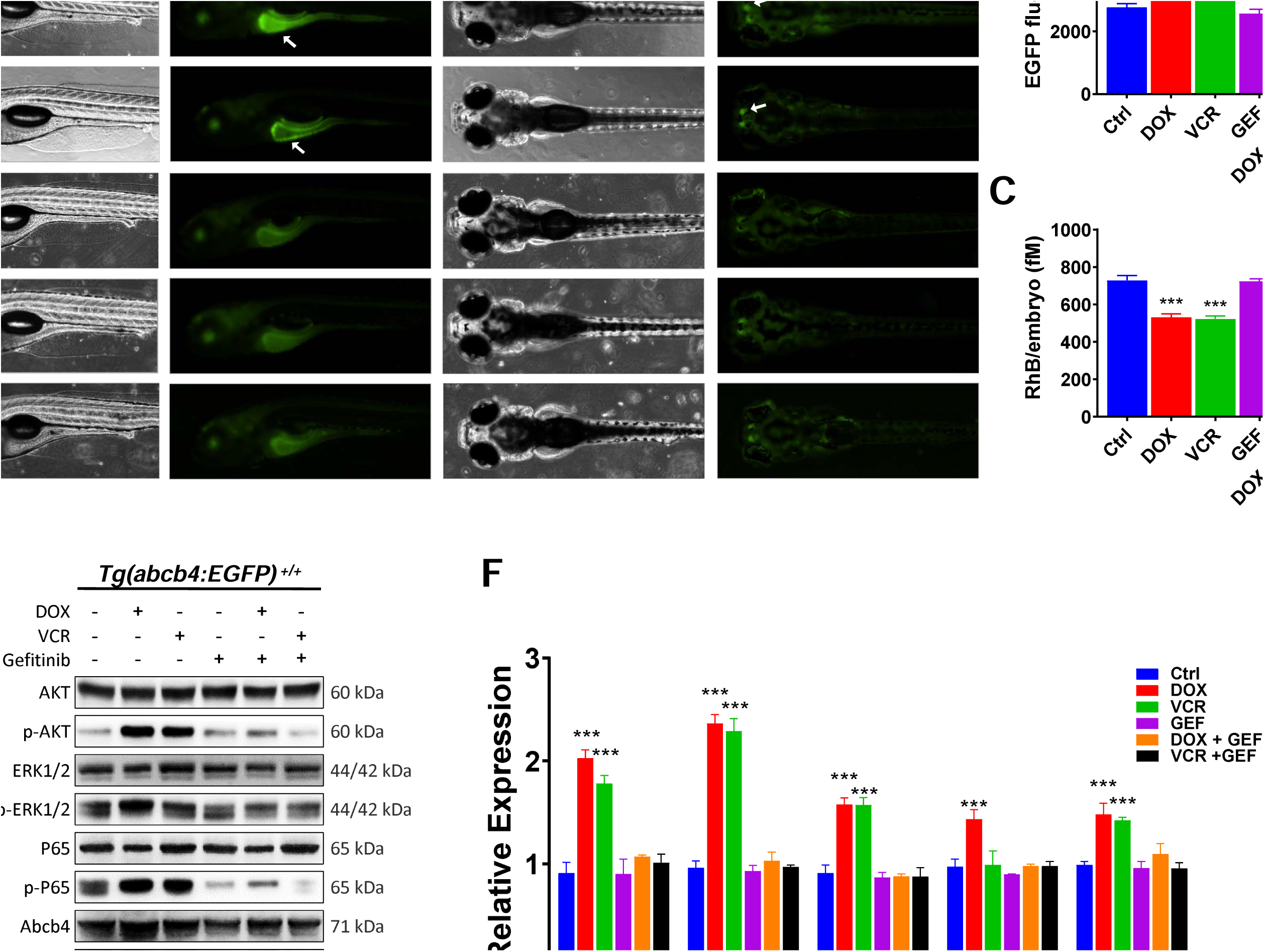
Effects of doxorubicin, vincristine, and gefitinib in *Tg(abcb4:EGFP)* zebrafish embryo. (A) Fluorescence micrographs of 5 dpf transgenic zebrafish embryos after treatment. The DOX and VCR group show the brighter fluorescence signal appears in the brain and intestine (white arrow) (B) Quantification of fluorescence intensity in transgenic zebrafish embryos. The results indicatet that DOX and VCR enhance fluorescence intensity, but the gefitinib inhibit the effect of DOX and VCR. (C) Quantification of rhodamine B (RhB) dye accumulation in 5 dpf zebrafish embryos when exposed to drugs. Low RhB dye accumulation indicates the enhanced ability of efflux in zebrafish embryos. (D) The level of AKT/p-AKT, ERK1/2, p-ERK1/2, P65/p-P65, abcb4, and EGFP were detected and the images were photographed. (F) The p-AKT, p-ERK1/2, p-P65, abcb4, and EGFP were analyzed by data normalization with β-actin antibody. Compared to control group, high level of p-AKT, p-ERK1/2, p-P65, abcb4, and EGFP presented in DOX and VCR groups, and there was no significant change in response to GEF treatment, DOX + GEF, and VCR + GEF groups. Data are presented as mean ± SD of three independent pools of 50 embryos. Statistically significant difference from controls was determined with one-way analysis of variance (ANOVA), followed by Dunnett’s test. *** *P*< 0.001 vs. control group. Ctrl, control; DOX, 15 μM doxorubicin; VCR, 25 μM vincristine; GEF, 15 μM gefitinib; DOX + GEF, 15 μM doxorubicin + 15 μM gefitinib; VCR + GEF, 25 μM vincristine + 15 μM gefitinib.

In addition, the potential regulatory effects of doxorubicin and vincristine on signaling pathways closely related to drug resistance were examined in *Tg(abcb4: EGFP)* transgenic zebrafish model. The embryos were treated with doxorubicin and vincristine alone or in combination with gefitinib. Western blotting assay also showed the increase in the expression of Abcb4 and EGFP after doxorubicin and vincristine treatment; and gefitinib normalized this increase (Figure 3D and F). Meanwhile, AKT, ERK, and P65 were activated upon the exposure of doxorubicin and vincristine, evident from the increase in phosphorylation level, whereas, gefitinib reversed this effect (Figure 3D and F). Taken together, these results demonstrate that Abcb4 induces drug resistance to doxorubicin and vincristine in the engineered *Tg(abcb4: EGFP)* transgenic zebrafish model, which is a suitable system for studying drug resistance and underlying mechanism.

### Abcb4-mediated drug resistance to doxorubicin and vincristine is ascribed to AKT/NF-κB and ERK/NF-κB signaling in transgenic zebrafish

Following the observation on the phenomena of drug resistance and alteration in the phosphorylation of AKT, ERK, and p65 signaling in the engineered *Tg(abcb4: EGFP)* transgenic zebrafish model, we further testified the underlying mechanism for the regulation of Abcb4-mediated drug resistance. LY294002 (AKT inhibitor), U0126 (ERK inhibitor), and BAY 11-7082 (NF-κB inhibitor) were utilized for pre-incubation of embryos for 2 hours. The embryos were treated with doxorubicin and vincristine alone, in which EGFP signal was increased and RhB accumulation was decreased as above observed (Figure 4A, B and C). Treatment of embryos with doxorubicin and vincristine combined with LY294002 or U0126 did not reverse these phenomena (Figure 4A, B and C); whereas combinatorial treatment of embryos with doxorubicin and vincristine and NF-κB inhibitor or LY294002 and U0126 together could normalized the inducing effects of doxorubicin and vincristine (Figure 4A, B and C). It suggests that NF-κB signaling could be the node for the regulatory machinery with the involvement of AKT and ERK signaling. To further demonstrate the participation of signaling pathways, we directly tested the phosphorylation of AKT, ERK, and p65 as the readout for the activation of AKT, ERK, and NF-κB pathways, respectively. The expression level of abcb4 and EGFP was significant induced after doxorubicin and vincristine treatment (Figure 5A, B, C, and D). Pre-incubation of AKT inhibitor did not completely reverse the inducing effect of doxorubicin or vincristine in embryos (Figure 5A, B, C, and D); similarly, pre-incubation of ERK inhibitor partially reverse the inducing effect of doxorubicin or vincristine in embryos (Figure 5A, B, C, and D). However, pre-incubation of AKT and ERK inhibitor together could fully reverse the inducing effect of doxorubicin or vincristine (Figure 5A, B, C, and D). Notably, AKT inhibitor exerted more potent effect in the setting of doxorubicin treatment, and ERK inhibitor showed more potent effect in the setting of vincristine exposure. Furthermore, NF-κB inhibitor exhibited the same effect as combined AKT and ERK inhibitor (Figure 5E and F). Taken together, these results suggest that AKT and ERK signaling pathways contribute to the abcb4-mediated drug resistance, with the convergence of NF-κB node.

**Figure 4.**
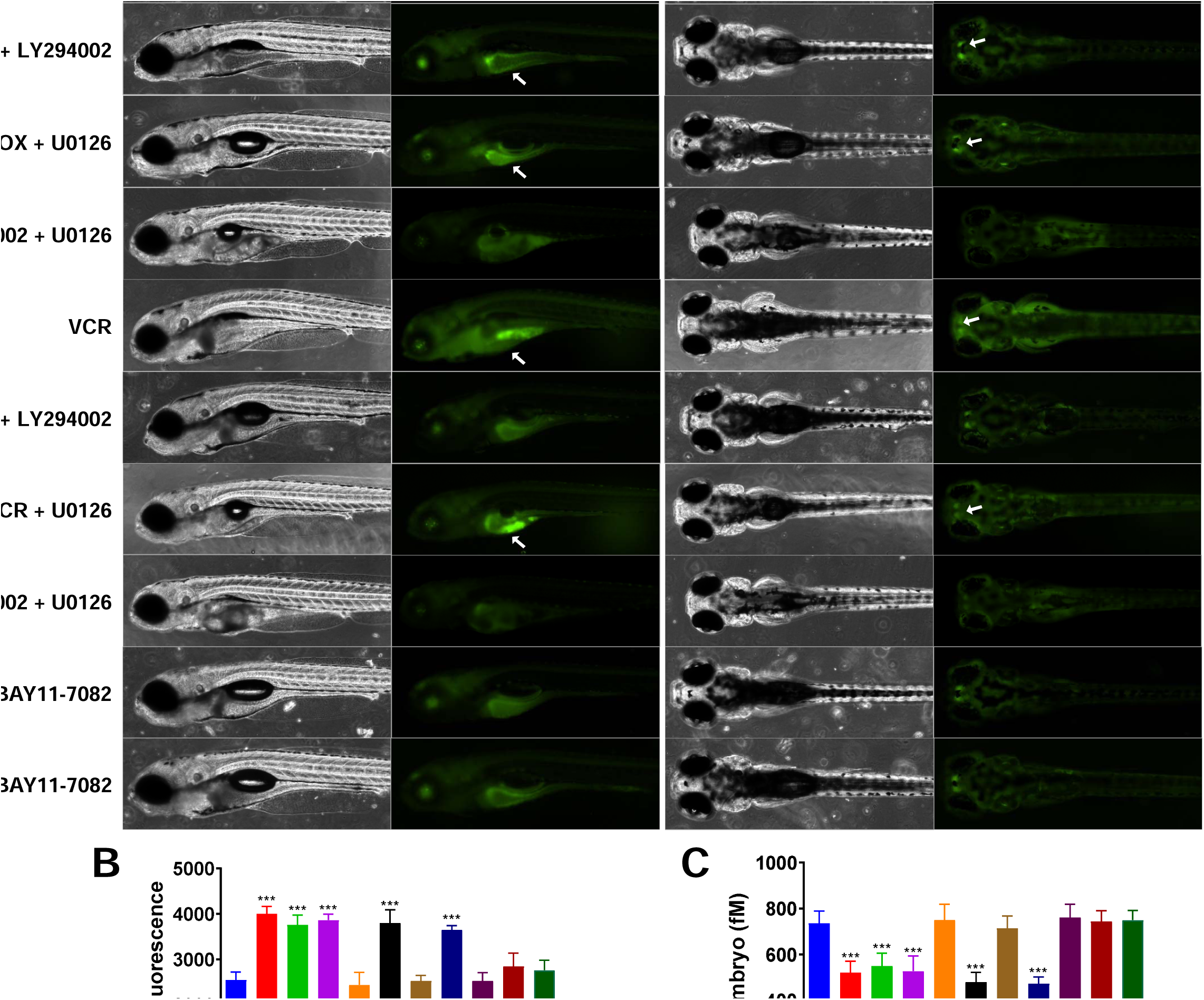
Effects of LY294002, U0126, and BAY 11-7082 in *Tg(abcb4:EGFP)* zebrafish embryo. (A) Fluorescence micrographs of 5 dpf transgenic zebrafish embryos after treatment. DOX, DOX + LY294002, DOX + U0126, VCR, and VCR + U0126 groups showed brighter fluorescence signal in the brain and intestine (white arrow). (B) Quantification of fluorescence intensity in transgenic zebrafish embryos. The results indicate that DOX, DOX + LY294002, DOX + U0126, VCR, and VCR + U0126 groups enhance fluorescence intensity, but the inhibitors suppress this enhancing effects. (C) Quantification of rhodamine B (RhB) dye accumulation in 5 dpf zebrafish embryos when exposed to various chemicals. Low RhB dye accumulation indicates the enhanced ability of efflux in zebrafish embryos. Data are presented as mean ± SD of three independent pools of 50 embryos. Statistically significant difference from controls was determined with one-way analysis of variance (ANOVA), followed by Dunnett’s test. *** *P*< 0.001 vs. control group. Ctrl, control; DOX, 15 μM doxorubicin; VCR, 25 μM vincristine; DOX + LY294002, 15 μM doxorubicin + 6.5 μM LY294002; DOX + U0126, 15 μM doxorubicin + 2.5 μM U0126; DOX + LY294002 + U0126, 15 μM doxorubicin + 6.5 μM LY294002 + 2.5 μM U0126; VCR + LY294002, 25 μM vincristine + 6.5 μM LY294002; VCR + U0126, 25 μM vincristine + 2.5 μM U0126; VCR + LY294002 + U0126, 25 μM vincristine + 6.5 μM LY294002 + 2.5 μM U0126; DOX + BAY 11-7082, 15 μM doxorubicin + 0.5 μM BAY 11-7082; VCR + BAY 11-7082, 25 μM vincristine + 0.5 μM BAY 11-7082.

**Figure 5.**
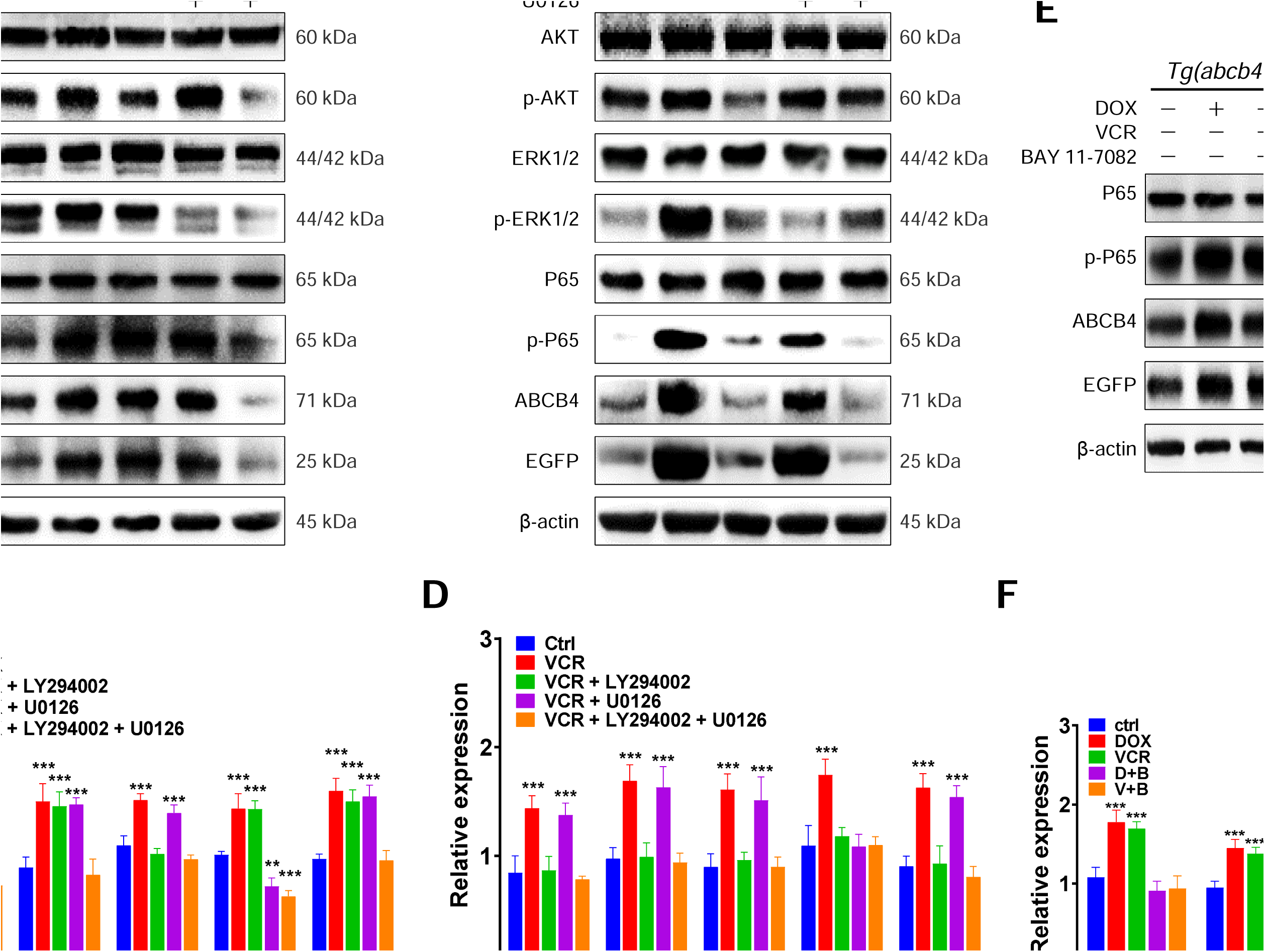
DOX and VCR regulate the expression of abcb4 through the AKT/NF-κB and ERK/NF-κB signaling pathways in *Tg(abcb4:EGFP)* zebrafish embryos. (A) The embryos were treated with 6.5 μM LY294002 and/or 2.5 μM U0126 inhibitor prior to 15 μM DOX, then the level of AKT/p-AKT, ERK1/2, p-ERK1/2, P65/p-P65, abcb4, and EGFP was detected and the images were photographed. (B) The level p-AKT, p-ERK1/2, p-P65, abcb4, and EGFP was analyzed by data normalization with β-actin antibody. Compared to control group, the high expression of abcb4 and EGFP presented in DOX, DOX+LY294002, and DOX+U0126 groups, whilst abcb4 was downregulated in DOX + LY294002 + U0126 group (* *P*< 0.05). (C) The embryos were treated with 6.5 μM LY294002 and/or 2.5 μM U0126 inhibitor prior to 25 μM VCR, then the level of AKT/p-AKT, ERK1/2, p-ERK1/2, P65/p-P65, abcb4, and EGFP was detected and the images were photographed. (D) The level p-AKT, p-ERK1/2, p-P65, abcb4, and EGFP was analyzed by data normalization with β-actin antibody. Compared to control group, increased expression of abcb4 and EGFP presented in VCR and VCR + U0126 groups, whereas there was no significant change in VCR + LY294002 and VCR + LY294002 + U0126 groups. (E) The embryos were treated with 0.5 μM BAY 11-7082 inhibitor prior to 15 μM DOX or 25 μM VCR, then the level of P65/p-P65, abcb4, and EGFP was detected and the images were photographed. (F) The p-P65, abcb4, and EGFP was analyzed by data normalization with β-actin antibody. Compared to control group, there was no significant change in DOX + BAY 11-7082 and VCR + BAY 11-7082 groups. Data are presented as mean ± SD of three independent pools of 50 embryos. Statistically significant differences from controls were determined with one-way analysis of variance (ANOVA), followed by Dunnett’s test. ** *P*< 0.01, *** *P*< 0.001 vs. control group.

## Discussion

In the present study, we engineered a transgenic zebrafish *Tg(abcb4:EGFP)*, in which the expression of EGFP is controlled by *abcb4* promoter. This model is expected to generate a new tool to study the mechanism of MDR in a different way from the manipulation of endogenous or exogenous ABCB1 expression [21, 22]. *Tg(abcb4:EGFP)* transgenic zebrafish has the capability of monitoring the transcriptional activity of *abcb4* promoter via visualizing EGFP signal, so as to estimate the transcriptional regulation of endogenous *abcb4* gene. This zebrafish model was assumed to facilitate the quick evaluation *in vivo* of MDR response through monitoring the level of EGFP protein fluorescence. Both gene and protein sequence analysis of ABC family proteins in zebrafish showed that the *abcb4* gene is closest to the human ABCB1 gene, suggesting abcb4 protein plays a similar role to P-gp protein [23]. Importantly, the expression pattern of Abcb4 in zebrafish which is similar to ABCB1 gene, mainly distributed in brain and intestinal tissues [24]. In this study, the same observation was found. We showed again this result by in situ hybridization examination with fluorescence labelling, which further highlights the potential application of this model in drug resistance screening for clinical use.

We evaluated the function of MDR/MXR response in zebrafish to estimate the similarity between abcb4 gene activation and P-gp gene activation. Therefore, the cells derived from engineered *Tg(abcb4:EGFP)* transgenic zebrafish was exposed to cytotoxic drug doxorubicin, vincristine because these two anticancer drugs were well-known P-gp substrates to induce P-gp gene expression in human cells and the drug responses are characterized. Previous studies showed that these three drugs have differential mechanism of action in cancer therapy. Clinically, the three drugs showed different degrees of MDR in the course of anti-tumor treatment [25, 26]. In human breast cancer and CML resistant cell lines, doxorubicin upregulates ABCB1 expression via PI3K/AKT/NF-κB and MAPK/ERK/NF-κB signaling pathways, so as to induce MDR response [27-30]. Our results showed that Tg(abcb4:EGFP) transgenic zebrafish gave a very similar intensity of the gene activation to those described by previous above mentioned studies with human cells. More interestingly, we also found that vincristine upregulated Abcb4 expression via AKT/NF-κB signaling pathway in zebrafish because the inhibitors of AKT/NF-κB signaling pathway blocked these response. AKT/NF-κB signaling pathway appeared to be a common mechanism because it has also been reported in studies on vincristine resistance in gastric cancer and myeloma [31, 32]. These results indicated that the abcb4 gene in zebrafish promoter was regulated in a similar way to the P-gp promoter in human. This further highlights that our engineered transgenic zebrafish can be used as an ideal animal model to study the MDR mechanism.

To further illustrate the potential of *Tg(abcb4:EGFP)* transgenic zebrafish model, we tested the abcb4:EGFP expression with cytostatic drug gefitinib, an EGFR inhibitor which binds to the catalytic domain of the RTK. Gefitinib has been shown to inhibit ABC transporter function and reverse ABCB1-mediated paclitaxel MDR in lung cancer cells and doxorubicin MDR in breast cancer cells, however, the underlying mechanism is still missing [33] [34]. We speculated that gefitinib may down-regulate the expression of Abcb4 through inhibiting the AKT/NF-κB and ERK/NF-κB signaling pathways, because the activation NF-κB has been shown to up-regulate the expression of ABCB1 in human [35, 36] and ABCB1 promoter contains NF-κB response element, which interacts with NF-κB and regulates the expression of *ABCB1* gene [37]. Effectively, in our *Tg(abcb4:EGFP)* transgenic zebrafish model, the addition of NF-κB inhibitor blocked the abcb4:EGFP expression. This result infers that abcb4 expression is controlled by NF-κB activaty upregulates and the cloned zebrafish *abcb4* promoter contains a conserved NF-κB response element, similar to ABCB1 promoter. Therefore, the experiment of EGFP expression in response to three drugs indicated that the transgenic zebrafish construction works correctly and it reflects the transcriptional activity of *abcb4* promoter which controls *EGFP* expression. This model can be considered reliable to intuitively and conveniently monitor the transcriptional activity of *abcb4* gene in zebrafish.

Our results are consistent with the expectation that abcb4 possesses functional properties of mammalian ABCB1, constituting an active barrier against chemical uptake and conferring resistance of embryos to ABCB1 substrates [24]. Meanwhile, transcriptional and functional MDR/MXR responses in zebrafish share overlapping features with the MDR/MXR mechanism in mammals, which is mainly reflected in the transcriptional regulation of *abcb4* gene by ABCB1 substrate and the effect of targeted agonists on the functional activity of abcb4 [38, 39]. Thusly, the present study provide a potential platform of MDR inducing activity for us to perform high-throughput drug screening for and MDR/MXR mechanism research.

## Acknowledgement

This project was supported in part by the National Natural Science Foundation of China (31860325), and in part by the Guizhou Province’s Science and Technology Major Project (Qian-P-Ren[2017]5611), and in part by the Guiyang’s Science and Technology Project (Zhukehetong[2017]5-1), and in part by the Non-profit Central Research Institute Fund of Chinese Academy of Medical Sciences(NO.2018PT31048, 2019PT310013).

## Supporting data

**Figure S1.**
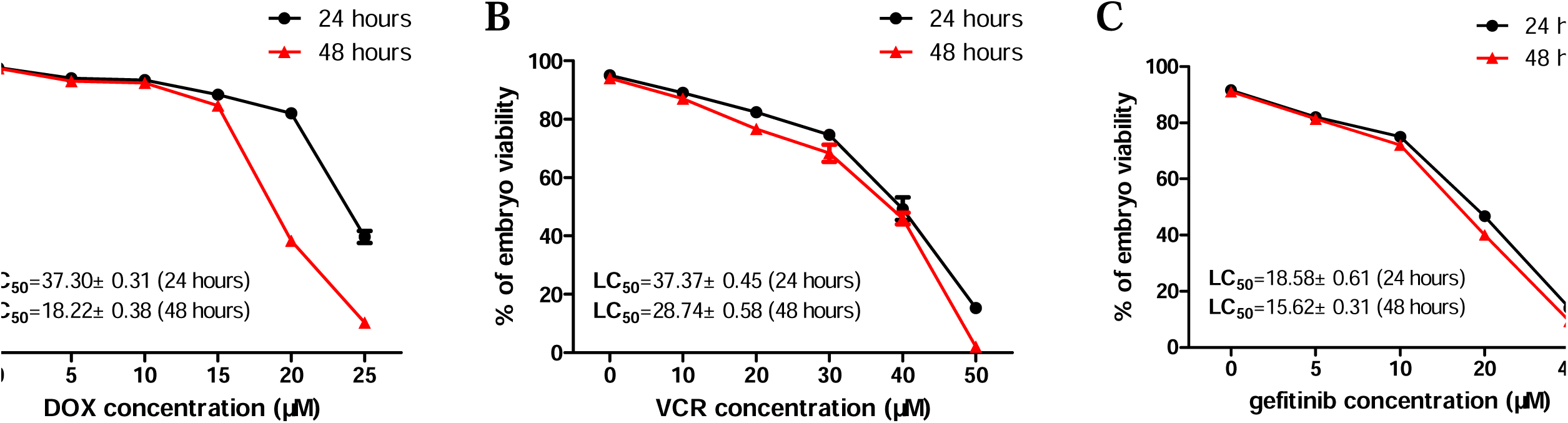
Calculated LC50 values with 95 % confidence intervals (CI) and the % differences of LC50s for zebrafish embryos after 24, 48 hpf exposures to different concentrations of doxorubicin (A), vincristine (B), and gefitinib (C).

