## Supplementary figures and images for "An engineered Abcb4 expressing model reveals the central role of NF-κB in the regulation of drug resistance in zebrafish"

### Supplemental Figure 1

#### Slide 1
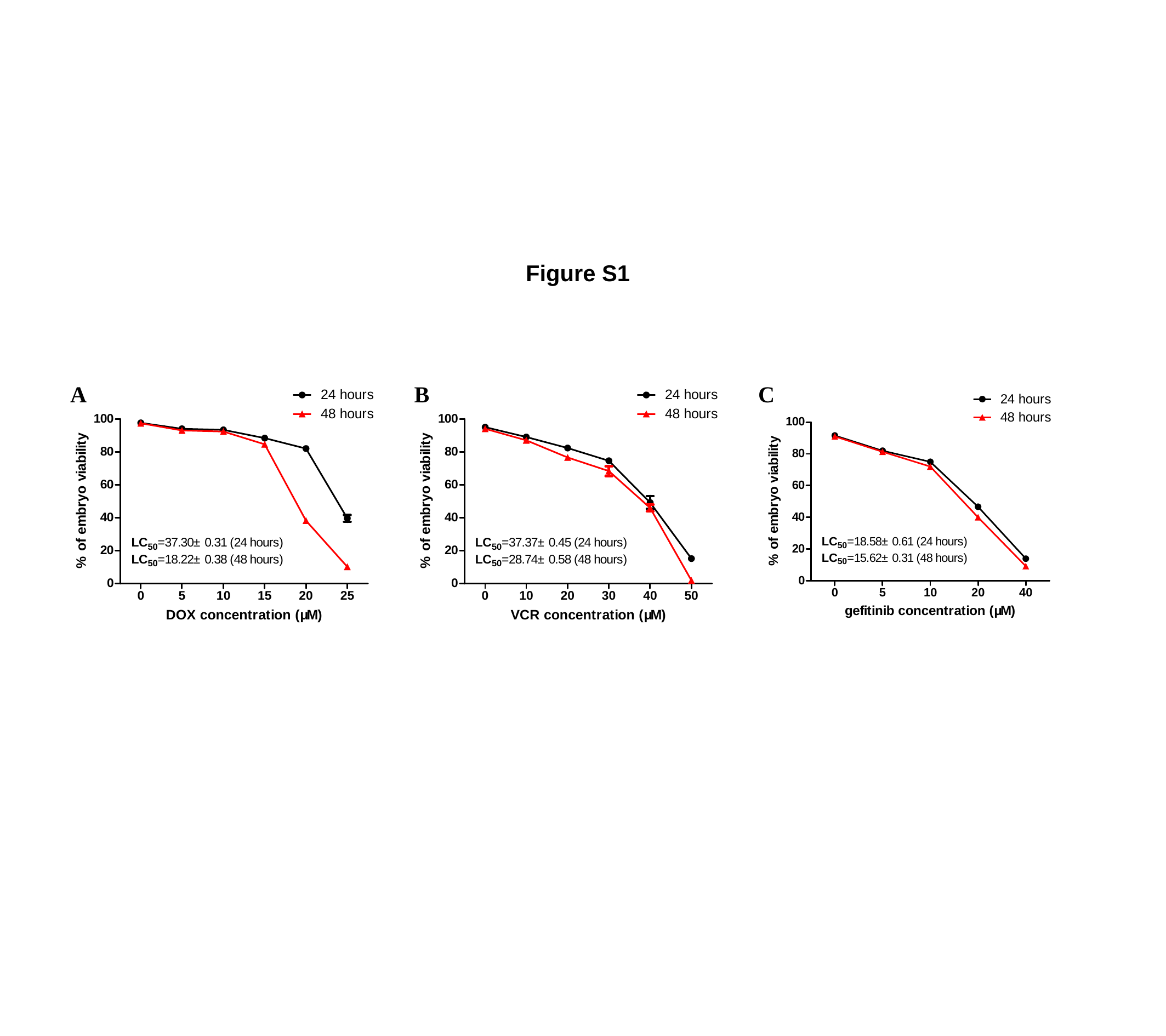

Figure S1
A
B
C
